# A multi-frequency whole-brain neural mass model with homeostatic feedback inhibition

**DOI:** 10.1101/2025.08.26.672269

**Authors:** Carlos Coronel-Oliveros, Fernando Lehue, Rubén Herzog, Iván Mindlin, Marilyn Gatica, Natalia Kowalczyk-Grębska, Vicente Medel, Josephine Cruzat, Raul Gonzalez-Gomez, Hernán Hernandez, Enzo Tagliazucchi, Pavel Prado, Patricio Orio, Agustín Ibáñez

## Abstract

Whole-brain models are valuable tools for understanding brain dynamics in health and disease by enabling the testing of causal mechanisms and identification of therapeutic targets through dynamic simulations. Among these models, biophysically inspired neural mass models have been widely used to simulate electrophysiological recordings, such as MEG and EEG. However, traditional models face limitations, including susceptibility to hyperexcitation, which constrains their ability to capture the full richness of neural dynamics. Here, we developed and characterized a new version of the Jansen-Rit neural mass model aimed at overcoming these limitations. Our model incorporates inhibitory synaptic plasticity (ISP), which adjusts inhibitory feedback onto pyramidal neurons to clamp their firing rates around a target value. Further, the model combined two subpopulations of neural cortical columns oscillating in α and γ, respectively, to generate a richer EEG power spectrum. We analyzed how different model parameters modulate oscillatory frequency and connectivity. We considered a model’s showcase, simultaneously fitting EEG and fMRI recordings during NREM sleep. Bifurcation analysis showed that ISP increases the parameters’ range in which the model exhibited sustained oscillations; the target firing rate acts as a bifurcation parameter, moving the system across the bifurcation point, producing different oscillatory regimes, from slower to faster. High frequency activity emerged from low global coupling, high firing rates, and a high proportion of γ versus α subpopulations. Importantly, ISP was necessary in the multi-frequency model to successfully fit EEG functional connectivity across frequency bands. Finally, ISP-controlled reductions in excitability reproduced both the slow-wave activity and the reduced connectivity in NREM sleep. Altogether, our model is compatible with biological evidence of the effects of E/I balance on modulating brain rhythms and connectivity, as observed in sleep, neurodegeneration, and chemical neuromodulation. This biophysical model with ISP provides a springboard for realistic brain simulations in health and disease.

**Author Summary:** Macroscale brain activity can be captured using techniques like EEG and fMRI. However, the granular or more detailed activity of neurons and neural masses is inaccessible. A solution is the use of whole-brain models, although they are not free from limitations, they can simulate EEG and fMRI recordings from mathematical equations and empirical data. One first limitation in these models is hyperexcitation. When the coupling between brain areas increases, brain areas might become aberrantly hyperexcitable if no compensatory mechanisms are considered. To address this, we introduce a mechanism in the model that dynamically modifies feedback inhibition to compensate for this excitability increase when running simulations. A second limitation is that many models fail to reproduce the spectral richness of EEG signals. EEG recordings reflect interweaving slower and faster rhythms, and some traditional models of EEG fail in capturing the spectral range of electrophysiological recordings. Here, we addressed this by combining two subpopulations of cortical columns within single brain areas, each one oscillating within the α and γ bands of EEG. Their combined activity generates EEG oscillations resembling the slower rhythms observed during sleep, and the faster ones triggered by increased attentional load. We ran different types of simulations and analyses to fully characterize our model. We observed that controlling system excitability is necessary to fully capture EEG connectivity and to simultaneously reproduce the EEG power spectrum and fMRI dynamics. Moreover, we showed that reduced/increased brain excitability is the cause of the emergence of the slowest/fastest EEG rhythms. The model can be used to characterize how connectivity and brain dynamics are altered in different types of conditions, such as chemical neuromodulation, drug delivery, altered states of consciousness, and neurodegenerative disorders. Our model is open access, well-documented, and introduced with tutorials, in the way to make it accessible to the whole neuroscience community.

## 1 Introduction

Understanding the biophysical mechanisms that give rise to large-scale brain dynamics remains a central topic in neuroscience [1, 2]. Generative models of brain activity offer a principled approach to bridge empirical observations and underlying mechanisms [3–5].

These models are mathematical architectures used to simulate neural population dynamics and are constrained by anatomical and neurophysiological priors. These models are particularly useful in capturing macro-scale patterns observed in functional magnetic resonance imaging (fMRI) and electroencephalography (EEG) [2, 6–8]. A key challenge, however, lies in setting these models to reflect biologically plausible excitation/inhibition (E/I) balances, especially when simulating whole-brain dynamics where hyperexcitability can dislocate realistic network behavior [6, 9, 10]. This hyperexcitability, manifested as increased excitatory inputs to neuronal populations, can stop or disrupt oscillatory activity in biophysical models. Another pitfall of some modeling architectures relies on their ability to generate realistic neurophysiological activity, such as the coexistence of slow and fast regimes of oscillatory dynamics [11]. Here, we build on a previously developed neural mass model used in dementia [7, 12] based on the Jansen-Rit equations [13, 14] and to introduce a biologically motivated inhibitory synaptic plasticity (ISP) mechanism [7, 9, 10] and multi-frequency behavior [11, 15]. Our approach allows simultaneous fitting of EEG and fMRI empirical data, providing a mechanistic account of neurophysiological changes during different brain states, i.e., wakefulness and NREM sleep.

Whole-brain models simulate the interactions of distributed brain regions and are commonly employed to explore how anatomical connectivity, neuromodulation, and local dynamics shape functional brain activity [2, 3, 5, 16]. These models vary in their level of abstraction. They range from pure abstract phenomenological models [17–19], to biophysically grounded models that aim to reproduce underlying neural principles [6, 20–22]. The latter can be used to propose biophysical mechanisms, particularly valuable in translational neuroscience, as they allow for proposing causal mechanisms in healthy and pathological brain dynamics [3, 7, 8, 23–26]. These models also facilitate the identification of biomarkers [7, 20, 27–29], and provide a framework for evaluating the potential effects of neuromodulation or pharmacological interventions in silico [7, 8, 30]. When constrained by empirical data, biophysical models can serve as testbeds for perturbational strategies aimed at restoring pathological brain dynamics [7, 30, 31].

Classical whole-brain models, for example, the Wilson–Cowan, Jansen–Rit and Dynamic Mean-Field models, remain the workhorses for simulating large-scale brain dynamics. These models often undergo bifurcations, such as Hopf bifurcations [32, 33], beyond which node dynamics may converge to stable fixed points and cease to oscillate. This transition disrupts oscillatory coherence and prevents the emergence of globally correlated network activity. Beyond excitability, empirical EEG signals span a broad frequency range, from δ to γ rhythms, each implicated in distinct cognitive and physiological processes. Capturing this spectral complexity is particularly important in clinical contexts such as dementia, where frequency-specific alterations in power and connectivity serve as functional signatures of neurodegeneration [7, 28, 34, 35].

Despite their utility, classical neural mass and mean field models face important limitations in simulating realistic brain-wide dynamics. One major gap arises from their susceptibility to hyperexcitation [6, 9, 10, 36], particularly when modeling large-scale networks with increased inter-regional coupling. Previous work has suggested that local regulatory mechanisms, such as ISP, can counteract hyperexcitability by dynamically adjusting inhibitory strength to maintain stable oscillations and network-wide integration [7, 9, 10]. The second gap relies on the failure of neural mass models in reproducing the spectral richness of human EEG signals [11]. These models, when using single frequency oscillators, might generate oscillations with narrow power spectra centered at the fundamental oscillatory frequency of the model. Our previous works have demonstrated that incorporating multiple subpopulations with different oscillatory frequencies within a cortical column can enhance spectral richness [7, 14, 15]. Nevertheless, only a limited number of models have been developed to date [10, 19, 24, 37–39] have attempted to jointly reproduce EEG- and fMRI-like brain dynamics, two complementary modalities that reflect different temporal and spatial scales of brain function.

To address these challenges, we characterize a biophysically inspired multi-frequency model, recently proposed in our earlier work [7, 15], which integrates ISP to stabilize network dynamics [9, 10]. We begin by analyzing the local effects of ISP through bifurcation analyses of single-node dynamics, demonstrating how inhibitory control modulates the transition between oscillatory and non-oscillatory states. We then explore how key model parameters influence the spectral content of simulated EEG signals. Using source-reconstructed EEG data from healthy participants, we fit the model to reproduce empirically observed functional connectivity (FC) across multiple frequency bands. Finally, we extend the model to simultaneously fit both EEG power spectra and fMRI FC during wakefulness and NREM sleep by modulating the E/I balance of neural masses. Through this work, we provide a generative framework, grounded in physiological mechanisms, that accounts for both spectral and connectivity features of brain dynamics.

## 2 Results

We used a modified version of the Jansen-Rit model (**Figure 1**) [7, 12]. The model combines empirical priors (**Figure 1A**) with a dynamical model for simulating neural activity from cortical columns (**Figure 1B**). Here, we provide a full characterization of the model from the single-node to the whole-brain dynamics. At the nodal level, our model consisted of two subpopulations of cortical columns (**Figure 1B**). The parameters of these subpopulations defined their intrinsic oscillatory frequencies. Specifically, the neural gains for excitatory, *A*, and inhibitory, *B*, populations, and the inverse of characteristic time constants for excitatory, *a*, and inhibitory, *b*, synapses were chosen to control the frequency of oscillations (**Table 2** with the model’s parameters). The parameters allow subpopulations to oscillate within the α or γ EEG canonical frequency bands [15]. The proportion of α versus γ neurons, *r*^α^, controls the contributions of each subpopulation to the EEG-like activity [7, 15], determining the spectral properties of individual brain areas. The other element of our model consists of ISP (**Figure 1B**), modeled as an additional differential equation for adjusting the feedback inhibition onto pyramidal neurons, *C*_4_, to dynamically set the firing rate of pyramidal neurons around a target value, *ρ*. A synaptic plasticity time constant, *τ* (the inverse of the learning rate), controls the speed of convergence to the target firing rate value.

**Figure 1.**
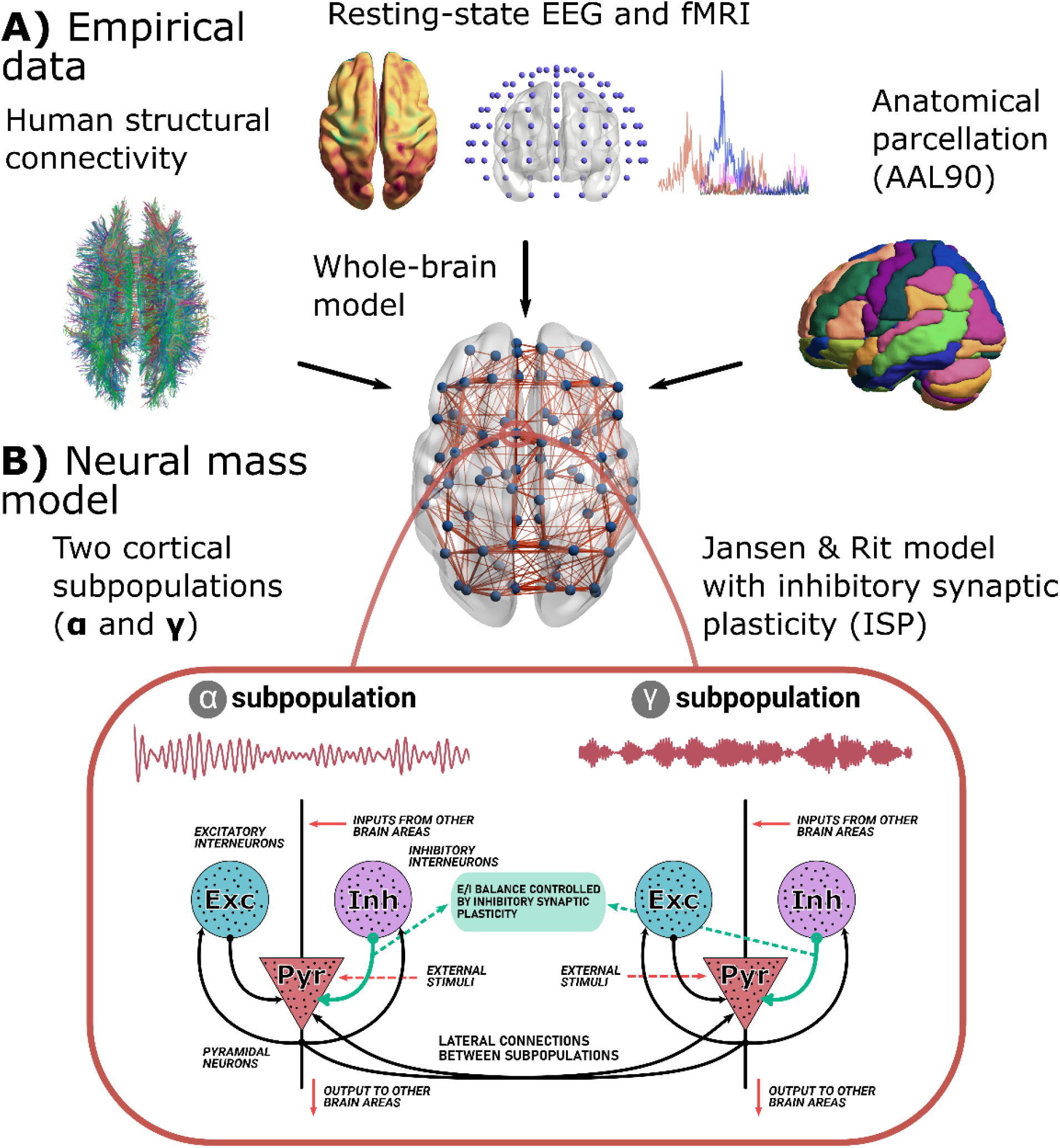
Whole brain neural mass model. **A)** The model is constrained with structural connectivity (SC, from DTI), fMRI/EEG functional connectivity (FC), and the EEG power spectrum. Brain areas were parcellated using the AAL90 brain parcellation. **B)** Each brain area in the whole brain model consisted of a modified version of the Jansen-Rit neural mass model. Each region is modeled using two subpopulations of neural masses with a frequency peak of oscillation within the α and γ bands of the EEG spectrum. Both subpopulations include the interactions between pyramidal and interneuron populations. The model also included inhibitory synaptic plasticity (ISP), as an additional equation to model the time courses of the feedback inhibition onto pyramidal populations, to reach a desired average firing rate. Figure partially modified from [7] with permission of the authors.

### 2.1 Analysis of Jansen-Rit single-node dynamics

We characterized the effect of plasticity by studying the model at the single-node level and containing only the α population (*r*^*α*^ = 1). Without plasticity, this corresponds to the original Jansen-Rit model in its most common implementation and parameter set (with a slight modification of neural gains and time constants, as presented in **Table 2**). **Figure 2A** shows a bifurcation diagram of this model against the *p* (external input) parameter, showing the attractors for the *x*_*0*_ variable (excitatory inputs to pyramidal neurons), the frequency of the detected oscillations, and two examples of output voltage. As it has been previously described [33, 40], the model presents robust oscillations of ~10Hz within a certain range of *p*.

**Figure 2.**
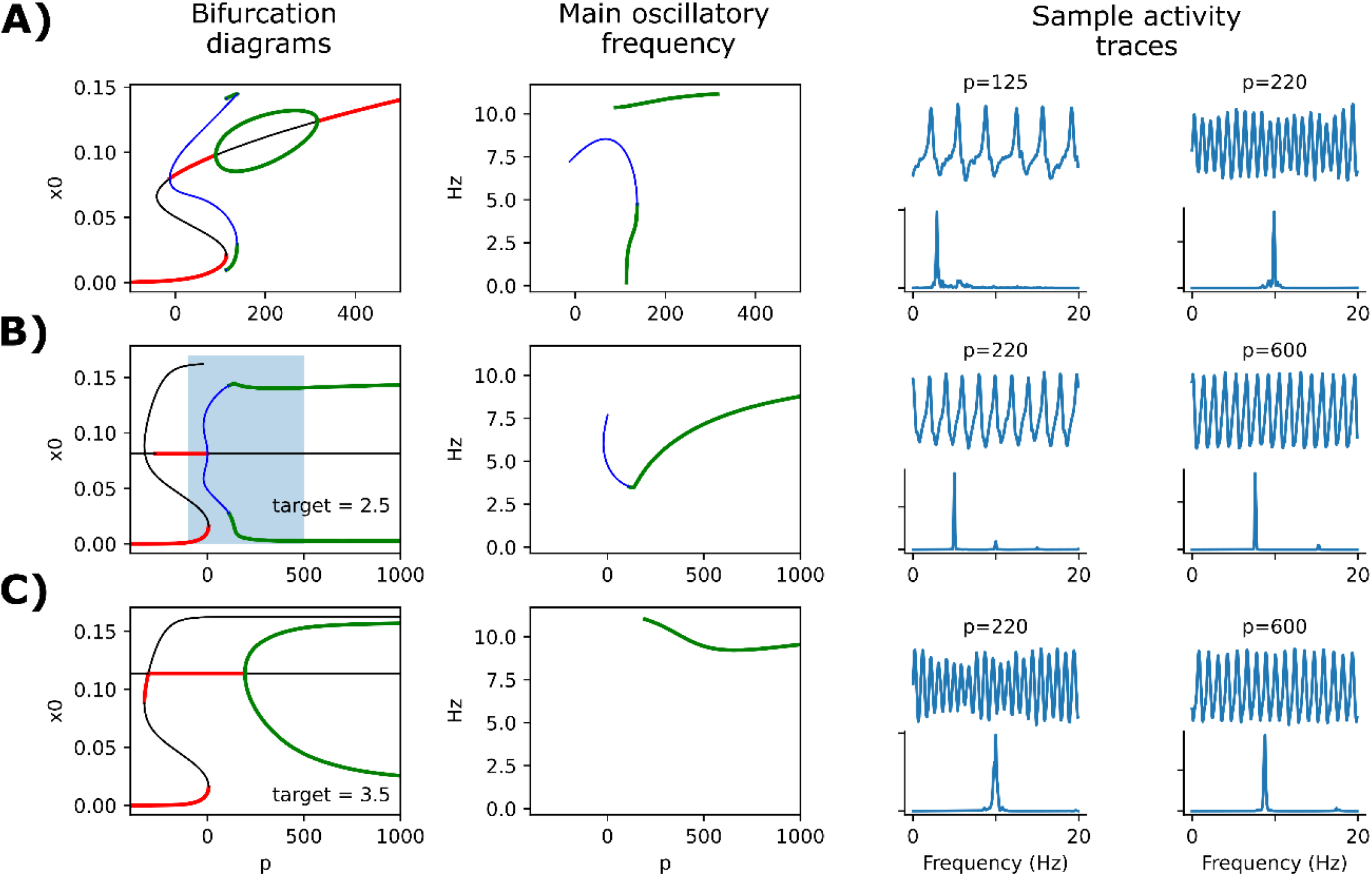
Bifurcation analysis of the model with and without plasticity. **A)** Bifurcation diagram, main oscillatory frequency, and two sample activity traces at the indicated *p* input values, for the original Jansen-Rit model without plasticity and consisting only of α (10 Hz) population. Below the traces, a power spectral density (PSD) plot is shown. In the left panel, red thick lines and black lines represent stable and unstable fixed points, respectively. Green and blue lines represent maxima and minima of stable (green) and unstable (blue) periodic orbits. The colors are maintained in the center panel to show the frequency (in Hz) of the associated oscillations. Simulations include a noise factor added to the *p* parameter. **B)** Same as in (A), with the addition of the homeostatic plasticity mechanism and a target for pyramidal firing rate *ρ* = 2.5 Hz. The shaded rectangle in the left panel denotes the range of *p* that is plotted in (A). **C)** Same as in (B), with the target *ρ* = 3.5 Hz.

The addition of homeostatic plasticity into the model dramatically reshapes the bifurcation diagram of the model (**Figure 2B-C**). Now, oscillations are sustained robustly in a much wider range of *p* (note the difference of scale in the *X* axes of **Figure 2B** vs **2A**), and the variables also oscillate more widely. The frequency of oscillation is not only dependent on the input *p*, but also on the target value *ρ* for the firing rate of the pyramidal population. When *ρ* is set to 3.5 Hz, oscillations are set very robustly around the α (10 Hz) band (**Figure 2C**). Also, with *p* = 220 Hz, the modulation of amplitude resembles very closely the behavior of the original model at the same input value [32, 33].

Then, we analyzed the effect of combining the two cortical subpopulations, α and γ, on nodal oscillatory frequency. We ran simulations without ISP to characterize the EEG power spectrum shape (**Figure 3A**). Using single-node simulations, we modified the *r*^α^ parameter to control the contribution of α versus γ neurons within the cortical columns. For *r*^α^ < 0.25, the single node oscillates within the γ EEG frequency band (> 30 Hz). On the other hand, *r*^α^ > 0.9 produces mainly α oscillations (with a frequency peak around 10 Hz). For intermediate values of *r*^α^, the model exhibits a richer power spectrum, with distributed power around all the possible canonical frequency bands. Using the multi-frequency model, we ran single-node simulations with *r*^α^ = 0.5 to check the model’s capability to clamp firing rates to target values. Across different values of the external stimulation, *p*, and target firing rates, ρ, the model can adjust the feedback inhibition (the inhibitory-to-excitatory local connectivity constant C4) to reduce/increase excitability and finally match the desired model firing rates (**Figure 3B-C**).

**Figure 3.**
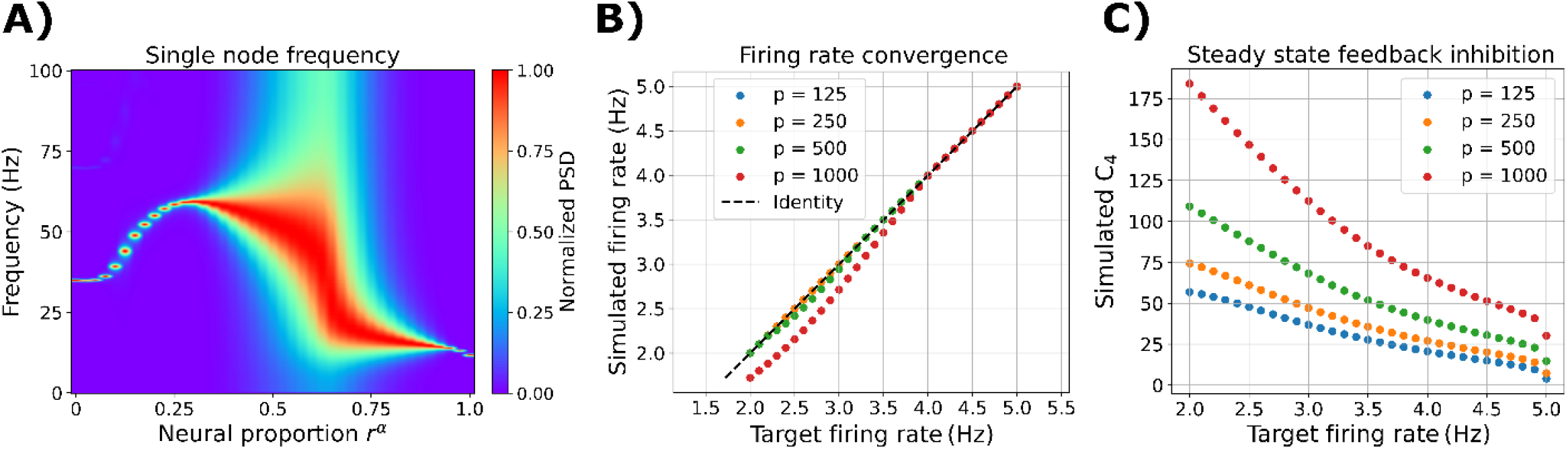
Single-node dynamic. **A)** Modulation of oscillatory frequency in single-node simulations. Normalized PSD as a function of α versus γ subpopulation proportion, *r*^α^, in the single-region model. The values shown here are the average of 50 random seeds. **B)** Noise-free simulations, with fixed *r*^α^ = 0.5, showing the target firing rates ρ versus the time-averaged simulated firing rates. **C)** Noise-free simulations, with fixed *r*^α^ = 0.5, showing the target firing rates ρ versus the time-averaged feedback inhibition, given by the time-inhibitory-to-excitatory local connectivity constant C_4_.

### 2.2 Whole-brain multi-frequency model with ISP

Here, we investigated the behavior of the multi-frequency model with ISP when many regions were coupled. In this setting, the ISP mechanisms were used to prevent hyperexcitation. Based on the results of the previous section, we first fixed the target firing rate *ρ* to 2.5 Hz and *r*^α^ = 0.5 [7, 41] (**Figure 4)**, as they generated robust oscillatory activity in a wide spectrum of frequencies. To connect different regions, we employed structural connectivity (SC) data of 45 healthy controls from the ReDLat consortium [42]. Data was parcellated into 90 regions using the AAL90 brain atlas [43] (**Table 1**), excluding all subcortical areas except the amygdala and hippocampus. We swept the global coupling parameter, *K*, from no coupling (*K* = 0) to strong coupling (*K* = 1) (**Figure 4A**). For *K* = 0, the local dynamics are constrained to solely γ oscillations. Increasing the global coupling to *K* = 1 generates α and θ rhythms. In the middle, with *K* = 0.5, a region of coexistence of faster (β, γ) and slower (θ, α) oscillations emerged at the whole-brain level. When fixing global coupling to *K* = 0.5, *r*^α^ = 0.5, and then manipulating target firing rates, *ρ*, we can shift brain dynamics from slower (for *ρ* < 2.5 Hz) to faster (for *ρ* > 2.5 Hz) regimes of activity (**Figure 4B**). Remarkably, we found no effects of the ISP time constant, *τ*, on the oscillatory frequency of the model (**Figure 4C)**.

**Table 1.** List of brain regions of AAL90 parcellation. The atlas comprises 90 cortical and subcortical brain areas (45 per hemisphere). Regions marked by * were included for fMRI simulations, but not for EEG.

| Brain Regions |  |  |  |
| --- | --- | --- | --- |
| Name | Abbreviation | Name | Abbreviation |
| Precentral gyrus | PreCG | Lingual gyrus | LING |
| Superior frontal gyrus (dorsolateral) | SFGdor | Superior occipital gyrus | SOG |
| Superior frontal gyrus (orbital) | ORBsup | Middle occipital gyrus | MOG |
| Middle frontal gyrus | MFG | Inferior occipital gyrus | IOG |
| Middle frontal gyrus (orbital) | ORBmid | Fusiform gyrus | FFG |
| Inferior frontal gyrus (opercular) | IFGoperc | Postcentral gyrus | PoCG |
| Inferior frontal gyrus (triangular) | IFGtriang | Superior parietal gyrus | SPG |
| Inferior frontal gyrus (orbital) | ORBinf | Inferior parietal gyrus | IPG |
| Rolandic operculum | ROL | Supramarginal gyrus | SMG |
| Supplementary motor area | SMA | Angular gyrus | ANG |
| Olfactory cortex | OLF | Precuneus | PCUN |
| Superior frontal gyrus (medial) | SFGmed | Paracentral lobule* | PCL |
| Superior frontal gyrus (medial orbital) | ORBsupmed | Caudate* | CAU |
| Rectus gyrus | REC | Putamen* | PUT |
| Insula | INS | Pallidum* | PAL |
| Anterior cingulate gyrus | ACG | Thalamus | THA |
| Median cingulate gyrus | MCG | Heschl gyrus | HES |
| Posterior cingulate gyrus | PCG | Superior temporal gyrus | STG |
| Hippocampus* | HIP | Temporal pole (superior) | TPOsup |
| Parahippocampal gyrus | PHG | Middle temporal gyrus | MTG |
| Amygdala* | AMYG | Temporal pole (middle) | TPOmid |
| Calcarine cortex | CAL | Inferior temporal gyrus | ITG |
| Cuneus | CUN |  |  |

**Table 2.** Model’s parameters.

| Parameter | Values | Units | Meaning |
| --- | --- | --- | --- |
| $A^\alpha$ | 3.9 | mV | Maximal amplitude of EPSPs ($\alpha$ population) |
| $A^\gamma$ | $32.5 a^\gamma / 1000$ | mV | Maximal amplitude of EPSPs ($\gamma$ population) |
| $B^\alpha$ | 26.4 | mV | Maximal amplitude of IPSPs ($\alpha$ population) |
| $B^\gamma$ | $440 b^\gamma / 1000$ | mV | Maximal amplitude of IPSPs ($\gamma$ population) |
| $a^\alpha$ | 120 | $\text{sec}^{-1}$ | Inverse characteristic time constant for EPSPs ($\alpha$ population) |
| $a^\gamma$ | 660 | $\text{sec}^{-1}$ | Inverse characteristic time constant for EPSPs ($\gamma$ population) |
| $b^\alpha$ | 60 | $\text{sec}^{-1}$ | Inverse characteristic time constant for IPSPs ($\alpha$ population) |
| $b^\gamma$ | 330 | $\text{sec}^{-1}$ | Inverse characteristic time constant for IPSPs ($\gamma$ population) |
| $r^\alpha$ | 0.5, [0,1] | Unitless | Proportion of $\alpha$ neurons |
| $\xi_{max}$ | 5 | Hz | Maximum firing rate of the sigmoid function |
| $r$ | 0.56 | $\text{mV}^{-1}$ | Slope of sigmoid function |
| $v_{th}$ | 6 | mV | Threshold of the sigmoid function |
| $M$ | [0,1] | Unitless | Structural connectivity matrix |
| $C$ | 135 | Unitless | Local connectivity |
| $C_1$ | C | Unitless | Excitatory-to-pyramidal neurons connectivity |
| $C_2$ | 0.8C | Unitless | Pyramidal-to-excitatory neurons connectivity |
| $C_3$ | 0.25C | Unitless | Inhibitory-to-pyramidal neurons connectivity |
| $C_4$ | 0.25C, variable | Unitless | Pyramidal-to-inhibitory neurons connectivity |
| $\langle p(t) \rangle$ | 220 | Hz | External input mean |
| $\sigma_p$ | 31 | Hz | External input standard deviation |
| $\rho$ | 2.5, [2.0,3.0] | Hz | Target firing rate for plasticity |
| $\tau$ | 2, [0.01, 50] | S | Inverse of plasticity's learning rate |
| $\beta$ | 1 | Unitless | Bounding exponent of plasticity |
| $K$ | [0,3] | Unitless | Global coupling |
| $\Delta t$ | 1 | msec | Integration step |
| $t_{eq}$ | 10 | sec | Discarded simulation time |
| $t_{sim}$ | [180, 660] | sec | Total simulation time |
| $t_{max}$ | [120, 600] | sec | Final time length of simulations |

**Figure 4.**
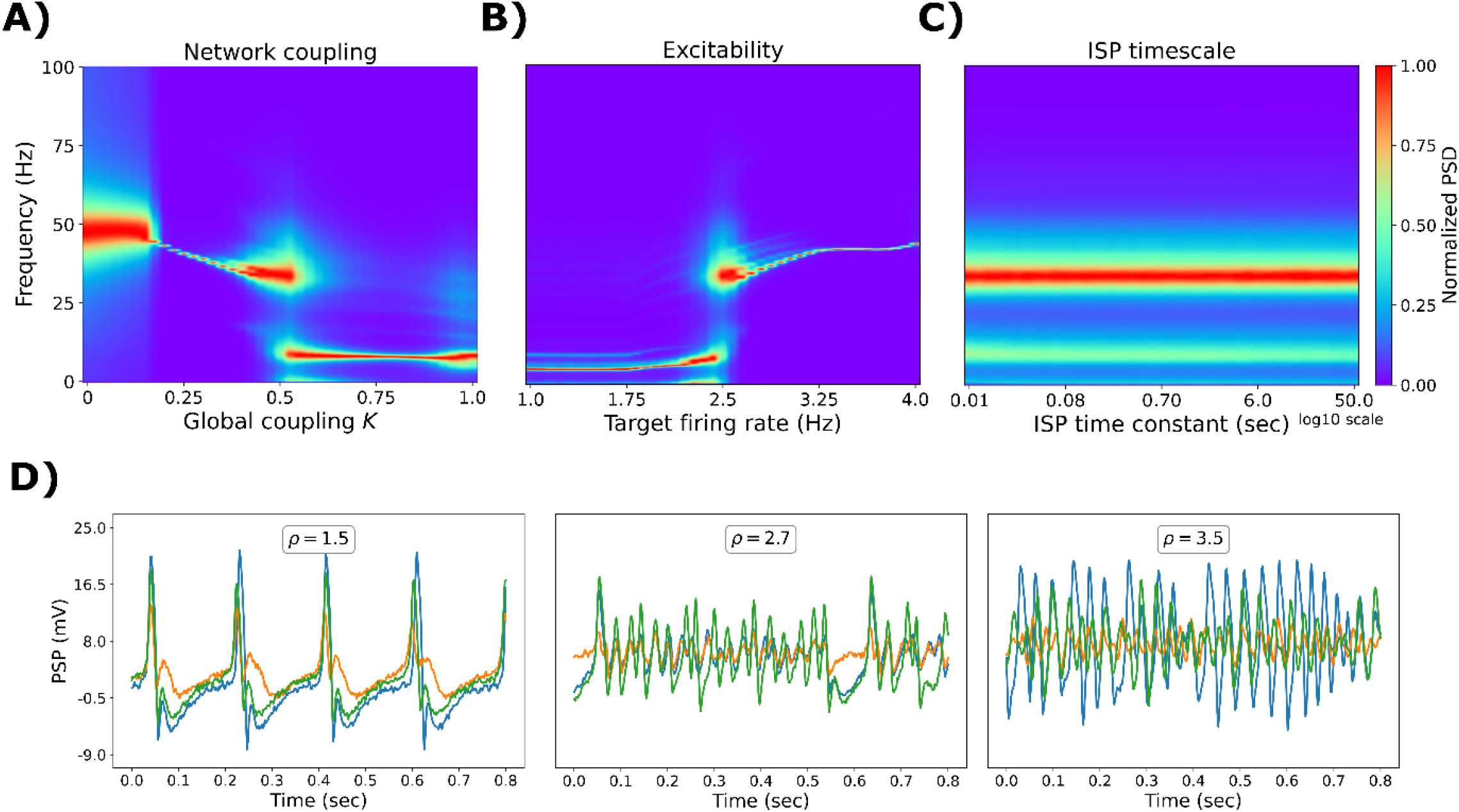
Modulation of oscillatory frequency in whole-brain simulations. Normalized power spectral density (PSD) as a function of: **A)** global coupling, *K*, in the whole-brain model, with a fixed target of *ρ* = 2.5 Hz. **B)** target (desire) firing rate in the whole-brain model, with a fixed *K* = 0.5. **C)** inhibitory synaptic plasticity (ISP) time constant, *τ*. **D)** Example of EEG traces for a fixed *K* = 0.5, *r*^α^ = 0.5, *τ* = 2 sec, and different values of *ρ*. The values shown here are the average of 50 random seeds.

In summary, we observed that slower oscillatory regimes emerge from a high proportion of α over γ neurons (*r*^α^) high global coupling (*K)* and low *ρ*. Faster oscillations appear in the opposite direction. Example PSP traces are depicted in **Figure 4D**, for *K* = 0.5, and *r*^α^ = 0.5. There, low *ρ* values produce δ-θ oscillations, intermediate values α rhythms, and the highest ones the faster β-γ neural activity.

### 2.3 Fitting to EEG source functional connectivity data

To further validate our model, we assessed its capacity to fit FC across the whole EEG power spectrum (**Figure 5**). We used the same SC data [42] and brain parcellation [43] described in the previous section, and the sourced EEG time series from the same participants [42]. We filtered the simulated and empirical EEG signals in the canonical frequency bands (δ, θ, α, and β). We calculated the signals’ envelopes, which were high-pass filtered (cut-off frequency of 0.5 Hz), and FCs were computed using the amplitude envelope correlation [44]. We compared the model without (**Figure 5A**) and with (**Figure 5B**) ISP. The *r*^α^ and *K* parameters were modulated to find the best combinations to reproduce EEG FC. Simulated and empirical matrices were contrasted using the Structural Similarity Index (SSIM, with SSIM = 1 being a perfect fit, 0 being the opposite) [30]. We aimed to simultaneously maximize SSIM for all EEG frequency bands (maximizing the average SSIM across bands).

**Figure 5.**
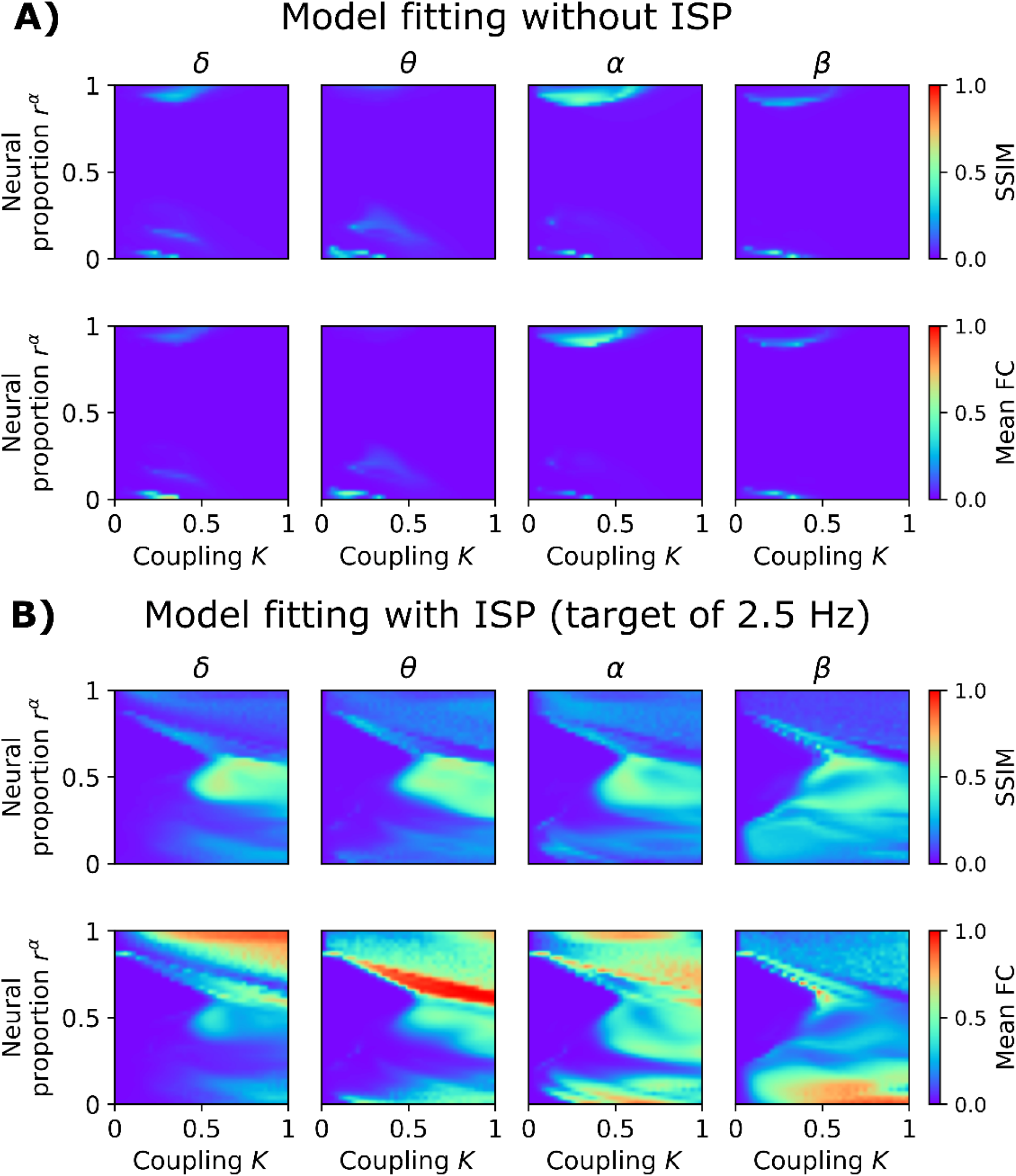
Goodness of fit of the model to empirical EEG FC. **A)** Model without inhibitory synaptic plasticity (ISP). The first row shows the structural similarity index (SSIM; captures the goodness of fit), and the second row represents the mean EEG FC of simulations. The axes consisted of the global coupling parameter, *K*, and the proportion of α versus γ subpopulations, *r*^α^. A value of SSIM = 1 indicates a good fit of the model. **B)** Goodness of fit using the model with ISP, and a fixed target firing rate, *ρ*, of 2.5 Hz. The values shown here are the average of 50 random seeds.

The model without ISP is characterized by a parameter space with a small region of correlated activity (**Figure 5A**). The model is unable to reach correlated activity before being hyperexcited, as no compensatory mechanisms were introduced in the form of ISP. Further, it cannot generate correlated activity when mixing α and γ subpopulations in cortical columns; that is, near 0 mean FC for *r*^α^ values around 0.5 suggests model frustration (inability to generate correlated activity). Including ISP in the model, with *ρ* = 2.5 Hz, enriches the parameter landscape with a wider range of parameters where correlations can emerge, from hypo to hyperconnectivity patterns, including intermediate connectivity regimes of activity (**Figure 5B**). Interestingly, correlations for intermediate values of *r*^α^ are now viable. Increasing global coupling in this scenario produced multi-frequency EEG FC connectivity, with maximal SSIM values in the proximity of *r*^α^ = 0.5 and *K* = 0.5.

We used the optimal parameters to compare the single-frequency model (with no ISP, *K* = 0.425, *r*^α^ = 1, SSIM = 0.14 averaged across frequency bands) and the multi-frequency model (with ISP, *K* = 0.675, *r*^α^ = 0.675, *ρ* = 2.5 Hz, SSIM = 0.56 averaged across frequency bands) to reproduce EEG FC (**Figure 6**). The SSIM values between the two models are presented in **Figure 6A**. There, the multi-frequency model outperformed the classical single-frequency model across all frequency bands (|D| > 1.2, representing a huge effect size) (**Figure 6A**). Example FCs are presented for the single and multi-frequency models, including the empirical FC matrices (**Figure 6B**).

**Figure 6.**
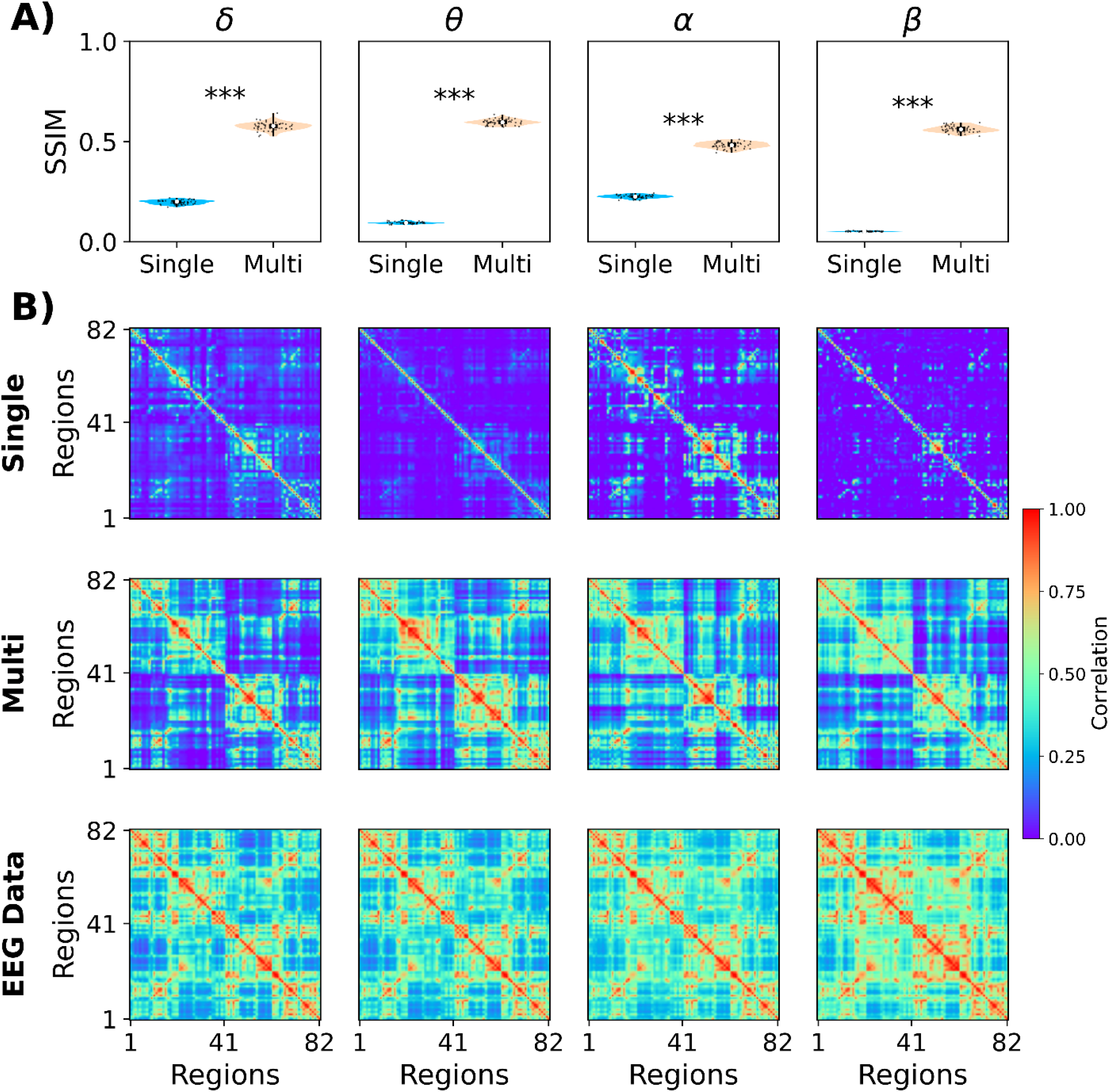
Comparison of the single- and multi-frequency models in fitting empirical EEG FC. **A)** The structural similarity index (SSIM) was used to measure the goodness of fit (SSIM = 1, perfect fit). The columns represent the different EEG frequency bands. **B)** Simulated EEG FCs from the single-frequency model. **C)** Simulated FCs from the multi-frequency model. **D)** Empirical EEG FC matrices. ***: |Cohen’s D| > 1.2. Each point corresponds to different models’ realizations (50 random seeds).

In this way, we demonstrated that ISP preserved E/I balance and enabled correlated activity without hyperexcitation. Also, the ISP mechanism combined with the multi-frequency model can generate multiband EEG FC across a wide range of parameter space.

### 2.4 Dual fitting to EEG and fMRI

As a final demonstration and validation of our model, we investigated its ability to jointly reproduce fMRI and EEG dynamics during NREM sleep. We used co-registers of EEG and fMRI during the transition from wakefulness to different stages of NREM sleep. fMRI recordings were collected from 71 individuals, with sleep stages labeled for each fMRI volume based on the simultaneously recorded polysomnographic data [45]. Here, we fitted our model to wakefulness and the N3 stage (deep sleep). We fixed *r*^α^ = 0.5, and systematically explored combinations of *K* and *ρ* drawn from a predefined parameter grid (define grid). We employed a connectome of young healthy participants from another study [46, 47]. We simulated EEG-like signals and then used a hemodynamic model [48] to transform the firing rates into fMRI BOLD-like signals. We aimed to fit two different observables, one from each recording modality. First, the empirical averaged EEG power spectrum at the sensor level (32 EEG channels), and second, the fMRI BOLD FC (considering the AAL90 brain parcellation). From the empirical and simulated power spectrum, we computed the θ, α, and β relative power, concatenated the values into a vector, and then compared the vectors using Clarkson’s distance [49]. So, we attempted to reproduce, using whole-brain simulations, the relative contribution of θ, α, and β oscillations in EEG. For fMRI BOLD FC, we compared simulated and empirical matrices using the SSIM.

The results, presented in **Figure 7**, suggest that solely using fMRI FC generates regions of the parameter space with model degeneracy in terms of goodness of fit (SSIM). That is, we observed dissimilar combinations of parameters with the same goodness of fit, regardless of whether the fitting was focused on wakefulness or NREM sleep. However, EEG fitting can be used as a mask to delimit the possible combinations of parameters that might reproduce empirical FC. In **Figure 7**, 1 – Clarkson’s distance represents the goodness of fit for simulated EEG, with values closer to 1 corresponding to a perfect fit to the empirical EEG relative power across frequency bands. Using a threshold of 0.85 for EEG fitting, we have a delimited parameter space where just a few parameters can simultaneously generate EEG and fMRI dynamics that closely match empirical data. For wakefulness, the best fit was found for higher coupling and medium firing rates (*K* = 1.44, *ρ* = 2.56 Hz, SSIM = 0.34, **Figure 7A**). In contrast, N3 sleep was better characterized by low coupling and low firing rates (*K* = 0.52, *ρ* = 2.1 Hz, SSIM = 0.29, **Figure 7B**).

**Figure 7.**
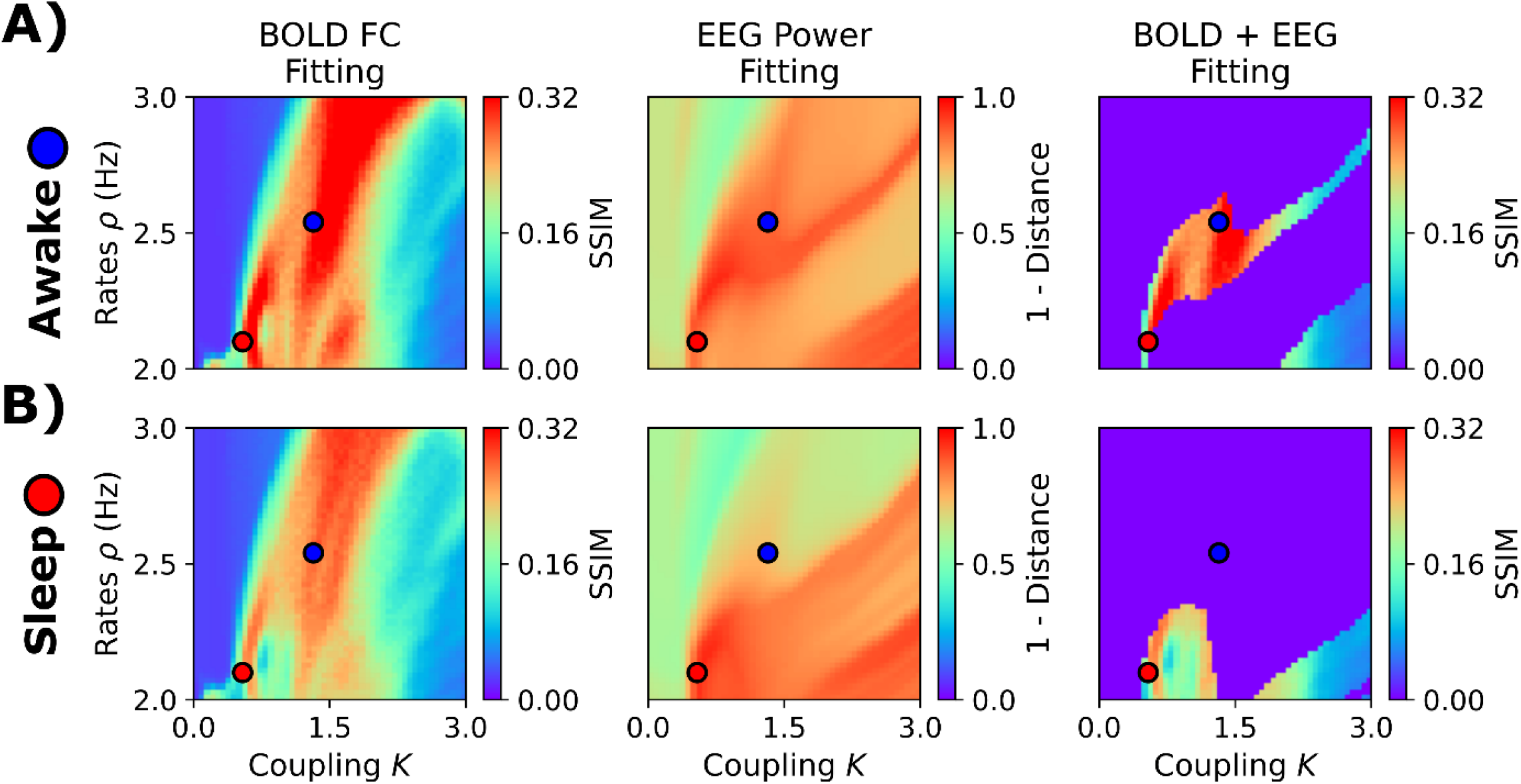
Fitting the EEG and fMRI sleep dataset. **A)** Fitting to awake condition. The first column corresponds to fMRI FC fitting, assessed using the structural similarity index (SSIM). The second column shows the goodness of fit to the EEG power spectrum. The last one corresponds to fMRI FC fitting using a threshold of EEG power fitting of 0.85. **B)** Fitting to EEG and fMRI NREM stage 3 (N3) data. The blue and red dots represent the best parameters (highest SSIMs when applying the EEG-fitting mask) for wakefulness (*K* = 1.44, *ρ* = 2.56 Hz) and N3 (*K* = 0.52, *ρ* = 2.1 Hz), respectively. The values shown here are the average of 50 random seeds.

Simulation outcomes are displayed in **Figure 8**. The simulated (**Figure 8A**) and empirical (**Figure 8B**) FC matrices showed decreased connectivity strength in N3 sleep. Simulated EEG-like signals showed high amplitude slow oscillations for N3 compared to wakefulness (**Figure 8C**). Finally, N3 and wakefulness simulated power spectra, when compared, presented the typical characteristics of the awake condition (peak in α) and deep sleep (power concentrated in the lowest frequency bands).

**Figure 8.**
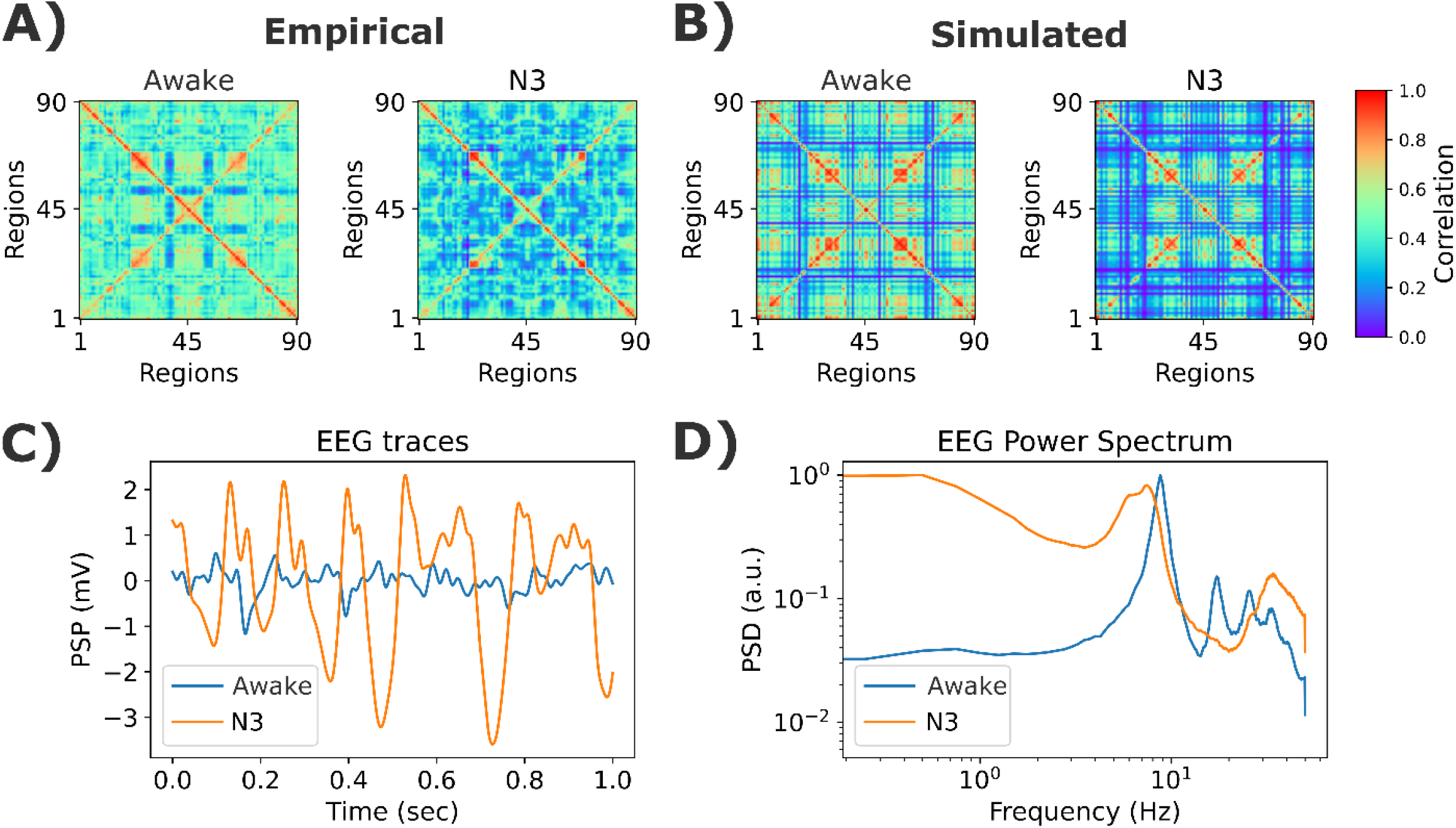
Example simulations for the EEG and fMRI sleep dataset. **A)** Empirical fMRI FC matrices for wakefulness and N3 stages. **B)** Best simulated matrices. **C)** Examples of EEG time courses from simulations. **D)** Simulated EEG power spectrum. The model’s parameters were *K* = 1.44 and *ρ* = 2.56 Hz, for wakefulness, and *K* = 0.52, *ρ* = 2.1 Hz, for N3 sleep. The values shown here are the average of 50 random seeds.

## 3. Discussion

In this work, we developed, characterized, and validated a semi-empirical Jansen-Rit whole-brain model able to simulate both EEG and fMRI BOLD dynamics from biologically plausible mechanisms. This model constitutes an improvement compared to the classical Jansen-Rit model in generating a richer power spectrum, allowing us to cover electrophysiological activity in the whole EEG frequency range. Further, the ISP control loop included in our model can be used as a mechanism for testing alterations in E/I balance associated with neuromodulation, drug delivery, and brain disorders. Using this version of the Jansen-Rit model, we simultaneously reproduced the EEG power spectrum and fMRI FC during wakefulness and sleep. In this way, we provided the computational neuroscience community with a model for whole-brain simulations with applications in basic and clinical neuroscience.

Our model extends the classical Janse-Rit model in two different ways. The first one is combining two distinct neural subpopulations to produce EEG signals with richer spectral content. Previous work [11] explored this approach using phenomenological models. Here, we implemented this multi-frequency mechanism within biologically inspired neural mass models. The rationale is grounded in the organization of cortical layers, where different neuronal subpopulations exhibit distinct input-output dynamics and generate frequency-specific rhythms [50]. Superficial layers tend to produce faster oscillations such as α and θ, while deeper and granular layers are associated with slower β and γ activity [51]. In our model, the EEG reflects a weighted summation of these laminar contributions, and the resulting broadband spectrum emerges from the combined activity of spatially and synaptically distinct rhythm-generating circuits. Bifurcation analyses of extended Jansen-Rit models show that under varying extrinsic inputs and dendritic time constants, a single cortical column can transition between oscillatory regimes spanning from δ to γ frequencies [32]. These transitions emerge from coexisting limit cycles and reflect biologically plausible shifts in input conditions, providing a structured framework for capturing the spectral diversity observed in empirical M/EEG.

The second addition of our model is the incorporation of ISP to modulate excitability and control the oscillatory regime of each subpopulation. In our framework, the target firing rate acts as a bifurcation control parameter [10]. By adjusting this parameter, the system can be placed either to the left or right of a Hopf bifurcation. The model produces slow oscillations for low firing rates, such as δ and θ. As the firing rate increases, the system transitions toward α oscillations. In the two-subpopulation version of the model, each subpopulation has its own limit cycle, and the nodal oscillatory activity becomes a combination of α and γ gamma rhythms. The balance between these rhythms depends on the target firing rate: higher values make γ activity more likely to be present, while lower values increase the α activity dominance. For very low firing rates, the model predominantly generates slow activity in the δ-θ range. Our model is compatible with other computational models that have shown how the modulation of excitability, driven by chemical neuromodulation [52–54] or pathophysiological processes [7, 8, 29], can produce shifts between different regimes, slower and faster, of brain dynamics. For example, [52] showed that an increase in neural adaptation, mediated by reduced acetylcholine [55], can generate slow waves in whole-brain simulations like the ones observed in NREM sleep. Other works [53] modeled sleep-like dynamics, diminishing excitatory feedback, and increasing global coupling. Similarly, our simulations showed that decreased excitability can generate EEG slow waves and realistic fMRI FC through ISP modulation. Other works using phenomenological models (Kuramoto oscillators with time delays), showed that faster oscillations emerge when using low global coupling, and, similar to our results, slower oscillatory regimes appeared when increasing inter-area connectivity [19].

By tuning model excitability, we were able to reproduce both EEG spectral slowing and reduced fMRI FC observed during NREM sleep [52–54]. These results are consistent with established neurophysiological findings, where NREM sleep is accompanied by decreased an overall decreased in neural excitability [56], triggered by an increased GABAergic inhibition and decreased levels of excitatory neuromodulators such as noradrenaline and acetylcholine [57, 58]. Our approach aligns with previous computational work modeling sleep-like dynamics through chemical neuromodulation, but here we demonstrate that a single mechanistic parameter (target excitability) is sufficient to reproduce multi-modal empirical data [52–54]. Beyond sleep, this model offers a framework to investigate pathological changes in excitability, such as those found in Alzheimer’s disease [7, 29, 59, 60]. Prior studies have used PET-based amyloid and tau deposition to modulate local excitability and mimic E/I imbalance [29, 61]; likewise, our model can potentially simulate shifts from early-stage hyperexcitability and hyperconnectivity toward later hypoexcitable, spectrally slowed states, reflecting the progression from preclinical to symptomatic stages of dementia [28, 62]. Finally, this framework can be extended to test the effects of pharmacological agents, or even non-invasive brain stimulation techniques, on cortical excitability and large-scale dynamics [7, 18, 30]. By capturing both EEG and fMRI features, our model provides a multimodal modeling tool for exploring how drugs or interventions may restore healthy brain function across diverse conditions and recording modalities.

This work has several limitations. First, excitability is modulated homogeneously across all brain regions, without accounting for known spatial differences in gene expression and neurotransmitter receptor distributions that influence local E/I balance [2, 23, 26, 29, 54]. A more realistic approach would involve modulating excitability in a region-specific manner based on empirical gene expression and receptor maps [3, 63]. Second, several mechanisms can modulate E/I balance, beyond ISP, that we did not consider in this work, i.e., changes in synaptic time constants or receptor kinetics directly [8, 32, 64]. Third, the model was validated on a relatively small cohort, which may limit generalizability. Larger and more diverse datasets are needed to assess robustness. Fourth, while the model requires further clinical validation, preliminary work from our team has shown good fit to empirical data in dementia [7, 12]. Fifth, the current implementation is limited to resting-state EEG and fMRI. Extending the model to simulate task-evoked activity, e.g., task-evoked β-γ changes, would help explore context-dependent changes in connectivity and cognitive processing [21, 23]. Finally, while the model is well-suited for testing brain stimulation and drug effects in theory, these applications remain to be validated experimentally.

Overall, we present a biologically grounded neural mass model capable of simulating realistic EEG and fMRI dynamics through a mechanistic modulation of excitability. The model bridges basic and clinical neuroscience, offering a framework to explore brain states, disease mechanisms, and interventions.

## 4 Methods

### 4.1 Participants and datasets

#### 4.1.1 EEG resting state and structural connectivity data (ReDLat)

We used data from 45 healthy participants from the ReDLat consortium [42]. Participants’ mean age was 71.2 ± 7.2 years (29 women). We considered both functional (FC) and SC data for this group. Full details can be found in our previous works [7, 65], but we briefly describe data acquisition and preprocessing below.

For EEG, participants were seated in an electromagnetically shielded EEG room, and instructed to stay still, awake, and with their eyes closed. As in previous works [7], resting-state EEG data were recorded for 10 minutes using a Biosemi ActiveTwo 128-channel system, with electrodes placed around the eyes to monitor blinks and movements. The signals were sampled at 1024 Hz and referenced to linked mastoids. Offline preprocessing included filtering (0.5 to 40 Hz), re-referencing all channels’ averages, and replacing malfunctioning channels using spherical interpolation [66]. Independent component analysis [67] and visual inspection was employed to correct blink artifacts and eye movements [68–71]. Brain sources were estimated using standardized low-resolution brain electromagnetic tomography (sLORETA) [72], resulting in time series data for 90 brain regions based on the AAL90 parcellation (**Table 1**). We only kept 82 brain areas from the AAL parcellation, including cortical regions plus the hippocampus and amygdala.

SC matrices were derived from diffusion tensor imaging (DTI) applied to diffusion-weighted imaging (DWI) data. Preprocessing was conducted using the FSL BEDPOSTX (Bayesian Estimation of Diffusion Parameters Obtained using Sampling Techniques) toolbox [73]. Post-preprocessing, each subject’s data yielded a 90 × 90 matrix representing connectivity between pairs of regions defined by the AAL parcellation. The final SC matrix was obtained by averaging the individual matrices across participants. We used the same 82 brain areas as in the EEG functional data.

#### 4.1.3 Structural connectivity data (connectomes of young participants)

We used SC data of healthy participants published in previous works [74]: 31 participants from the non-video game players group, mean age of 24.4 ± 3.0 years, all males. The SC data acquisition involved MR imaging on a 3-Tesla MRI scanner with a 32-channel phased array head coil. DTI data were preprocessed using the Pipeline for Analyzing Brain Diffusion Images software [75]. The processing steps included artifact correction, tensor construction, and registration to the MNI space, followed by parcellation using the AAL atlas. The 90 x 90 connectivity matrices were generated by calculating the number of fibers connecting each pair of brain regions, normalized between 0 and 1, and averaged across participants.

To improve model fitting, SCs connections were added in the anti-diagonal of the matrix to account for homotopic (inter-hemispheric) connections, which are typically underestimated using DTI [76]. Further details on data acquisition and preprocessing can be found in [75] and [77], and further information about participants and the original studies in [74] and [47].

#### 4.1.3 fMRI-EEG co-registering during wakefulness and NREM sleep

This dataset was originally used in [78].A total of 63 non-sleep-deprived subjects were scanned in the evening (starting from ~8:00 PM) and preliminarily included in the study (written informed consent; approval by the local ethics committee). A subset of 15 individuals with fMRI volumes in all W, N1, N2, and N3 stages was included in this study. All participants were scanned in the evening and instructed to close their eyes while lying still and relaxing. 1505 volumes of T2*-weighted echo-planar images were acquired using the following parameters: TR/TE = 2,080/30 msec, matrix 64 × 64, voxel size 3 × 3 × 2 mm^3^, distance factor 50%, field of view [FOV] 192 mm^2^ at 3 T (Siemens Trio). EEG recordings were simultaneously acquired with fMRI, using an EEG (modified BrainCapMR; Easycap) cap with 30 channels (sampling rate 5 kHz, low pass filter 250 Hz), along with a polysomnographic setup that included electromyography on the chin and tibia, electrocardiogram, bipolar electrooculography, and pulse oximetry. MRI artifact correction was performed using the average artifact subtraction. The fMRI data were realigned, normalized to MNI space, and spatially smoothed using Statistical Parametric Mapping (SPM8) with a Gaussian kernel (8 mm^3^ full width at half maximum). Sleep staging was performed by an expert according to the AASM criteria [79].

### 4.2 Functional connectivity estimation

For EEG, we filtered signals in the common EEG frequency bands: δ (0.5 – 4.0 Hz), θ (4.0 – 8.0 Hz), α (8.0 – 13.0 Hz), and β (13.0 – 30.0 Hz) using a 3rd-order passband Bessel filter. Then, we used the Hilbert transform to get the signals’ envelopes, then we filtered the envelopes with a 3rd-order Butterworth high-pass filter (cut-off frequency of 0.5 Hz), and finally, we built the FC by computing the Pearson’s correlation between all pairs of areas [44]0.

For fMRI, we filtered the signals between 0.01 – 0.08 Hz first using a 3rd-order Bessel filter, and then we built the FC matrix using Pearson’s correlation between all pairs of areas.

### 4.3 Power spectrum analyses

For both empirical and simulated data, the EEG power spectrum was calculated using the Welch method with 2-sec length time windows with 50% overlap [80]. The power was extracted for each EEG band and divided by the total power of the spectrum between 0.5 and 30 Hz.

### 4.4 Whole-brain model

Whole-brain activity at the source level was modeled using a modified version of the Jansen-Rit neural mass model [7, 13, 15]. In this model, each brain region consists of two subpopulations of neural masses, each tuned to oscillate in either the α (around 10 Hz) or γ (around 45 Hz) frequency bands of the EEG spectrum. The contribution of these subpopulations to the generation of the postsynaptic potential (PSP) of pyramidal neurons is weighted by the parameter *r*^*α*^, which reflects the proportion of *α* versus *γ* subpopulations within the brain areas, as indicated in [15]. Specifically, *r*^*α*^ = 1 indicates a full contribution from the *α* subpopulation, while *r*^*α*^ = 0 indicates none. Each subpopulation is composed of pyramidal neurons, along with excitatory and inhibitory interneurons. The PSPs, *ν*, are integrated and transformed into firing rates using a sigmoid function:

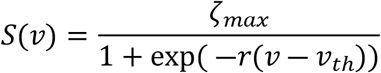

where *ζ*_*max*_ is the maximal firing rate output, *r* is the slope, and *ν*_*th*_ is the voltage threshold. The inverse operation is conducted by a PSP block, which convolves the firing rates of the neurons with an impulse response function defined as:

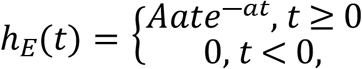

for the excitatory PSPs, and

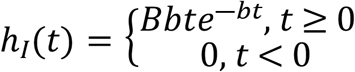

for inhibitory PSPs. Here, *A* (*B*) and *a* (*b*) correspond to the maximal amplitude and inverse characteristic time constant of the excitatory (inhibitory) PSPs, respectively. On a macroscopic level, brain areas *i* and *j* are connected using an SC matrix *M* derived from DTI data. The coupling strength was scaled by a global parameter *K* and, considering that long-range projections are primarily excitatory [81, 82], connections between brain regions involved only pyramidal neurons[7]. Each region received background input *p*(*t*), with values randomly sampled from a normal distribution with a mean ⟨*p*(*t*)⟩ = 220 Hz and a standard deviation σ_*p*_ = 31 Hz. The complete system of equations for the α subpopulation was:

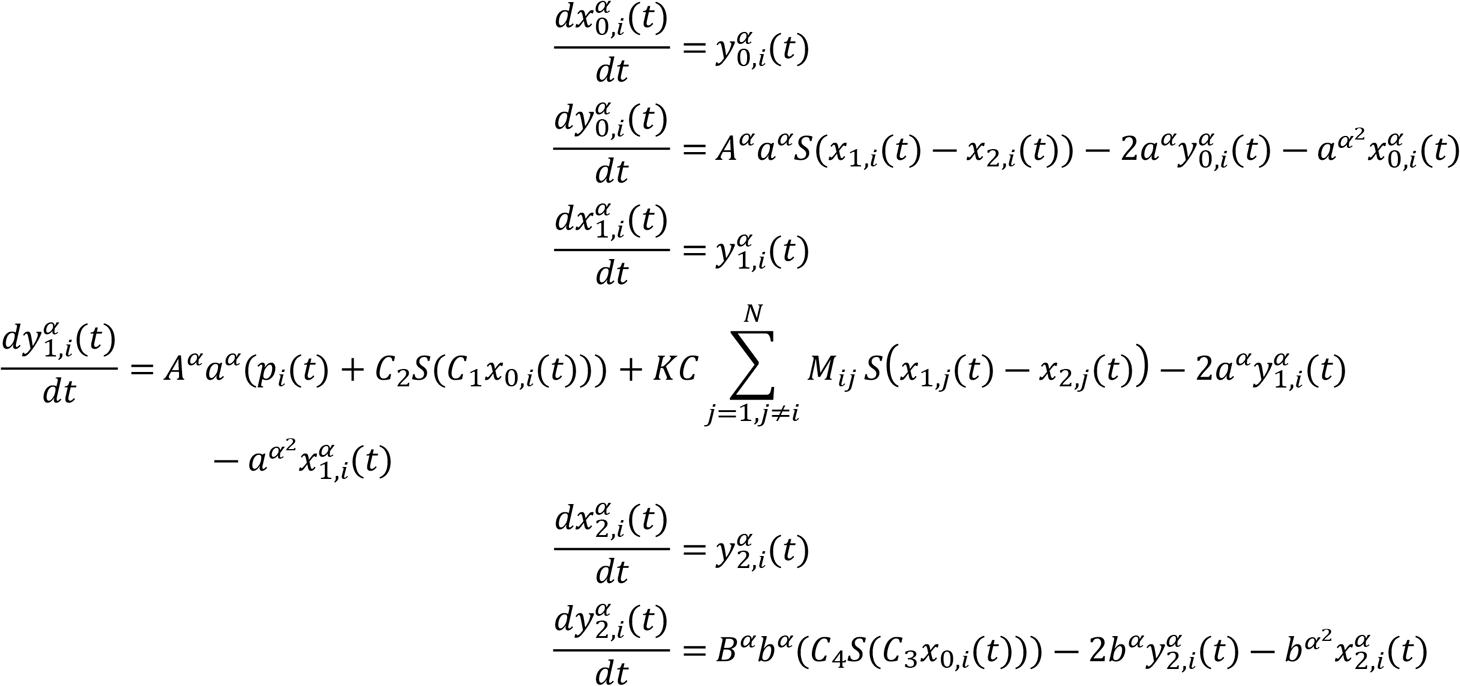

The first pair of equations model the excitatory feedback loop, the second pair models the outputs of pyramidal neurons, and the third represents the inhibitory feedback loop. Neuron populations were connected via constants *C*_*1*_, *C*_*2*_, *C*_*3*_ and *C*_*4*_, which are scaled by a local connectivity constant, *C*. Identical equations apply to the γ subpopulations, differing only by the γ superscript. The final output of the model produced EEG-like signals in source space, featuring a richer power spectrum. The model’s parameters are summarized in **Table 2**.

Additionally, following [9], we integrated inhibitory synaptic plasticity into the model. This mechanism controls the firing rate of pyramidal neurons, preventing hyperexcitation, which can occur when global coupling increases. Synaptic plasticity was introduced as an additional differential equation:

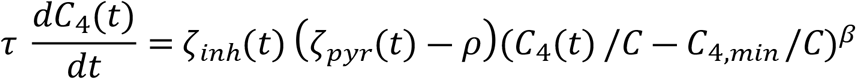

This equation dynamically updates feedback inhibition to regulate the firing rate of pyramidal neurons, thus preventing saturation of the sigmoid function (hyperexcitation). Here, *τ* represents the inverse learning rate, *ζ*_*inh*_ and *ζ*_*pyr*_ are the firing rates of inhibitory interneurons and pyramidal neurons at time *t*, respectively, *ρ* is the target firing rate, and β is a bounding exponent controlling convergence to *C*_*4,min*_. We used β = 1 (soft-bound), though other values are possible [83]. The key parameters for plasticity are the learning rate, *τ*, and the target firing rate, *ρ*, which were set as default to *τ* = 2 sec and *ρ* = 2.5 Hz [9, 41].

We solved the system of differential equations using the Euler-Maruyama method with a 1 msec integration step. The filtering and FC estimation procedures were the same as those used for the empirical signals.

### 4.5 Hemodynamic model

We simulated fMRI-like signals using the pyramidal firing rates *ζ*_*pyr,i*_(*t*) and a generalized hemodynamic model as presented by Stephan et al. [48]. In this model, an increase in the firing rate *ζ*_*pyr,i*_(*t*) initiates a vasodilatory response *s*_*i*_ leading to blood inflow *f*_*i*_, and subsequent changes in blood volume *v*_*i*_ and deoxyhemoglobin content *q*_*i*_. The system of differential equations governing this process is

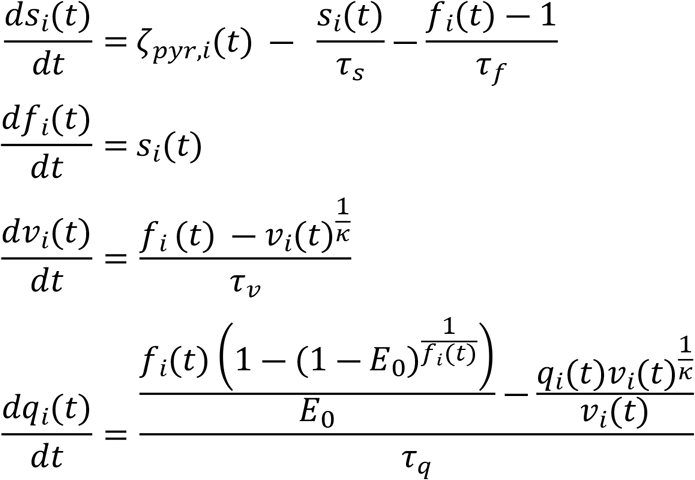

where the time constants *τ*_*s*_ = 0.65, *τ*_*f*_ = 0.41, *τ*_*v*_ = 0.98, and *τ*_*q*_ = 0.98 correspond to signal decay, blood inflow, blood volume, and deoxyhemoglobin content, respectively. The stiffness constant *κ* (resistance of the veins to blood flow) and the resting-state oxygen extraction rate *E*_*0*_ were set to *κ* = 0.32 and *E*_*0*_ = 0.4. The BOLD-like signal for the node *i*, denoted as *B*_*i*_(*t*), is a nonlinear function of *q*_*i*_(*t*) and *v*_*i*_(*t*):

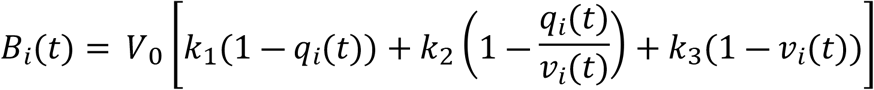

where *V*_*0*_ = 0.04 represents the fraction of deoxygenated venous blood at rest, and *k*_*1*_ = 2.77, *k*_*2*_ = 0.2, *k*_*3*_ = 0.5 are kinetic constants.

We solved the system of differential equations using the Euler method with a 10-msec integration step. The resulting fMRI-like signals were then down-sampled to match the empirical signal’s sampling rate (TR = 2.08 sec). The filtering and FC estimation followed the same procedures applied to the empirical signals.

### 4.6 Simulations

We first simulated the activity of an isolated node of the model (*K =* 0) without plasticity. There, we simulated 180 sec and discarded the first 60 sec (for a final length of 120 sec), downsampled the signals to 200 Hz sampling rate (to reduce memory consumption) and calculated the Normalized Power Spectrum (NPS, obtained using the Welch method and dividing by the maximum value of spectral density). Simulations were run against variations of *r*^*α*^ in 41 equidistant values between 0 and 1 (**Figure 3**).

We then ran a whole-brain simulation, where nodes represent brain areas that are connected using the ReDLat SC matrix. We simulated 660 sec, discarded the first 60 sec, and obtained the NPS (averaged across brain areas) against variations in different parameters (average of 50 seeds in **Figure 4**): we varied *K* over 41 equidistant values between 0 and 1 (upper right, fixing *r*^*α*^ = 0.5, *ρ* = 2.5 Hz, *τ* = 2 sec); *ρ* over 41 equidistant values between 1 and 4 Hz (lower left, fixing *r*^*α*^ = 0.5, *K* = 0.5, *τ* = 2 sec); and *τ* over 41 equidistant and log spaced values between 0.01 and 50 sec (lower right, fixing *r*^*α*^ = 0.5, *ρ* = 2.5 Hz, *K* = 0.5).

We fitted the model to the average FC matrices of healthy individuals in the ReDLat dataset [65], simulating 660 sec and discarding the first 60 sec of adaptation. We fixed *ρ* = 2.5 Hz and *τ* = 2 sec, and simultaneously varied the model’s global coupling *K* and *r*^*α*^ between 0 and 1 (41 values each), first without ISP (**Figure 5A**) and then with ISP (**Figure 5B**). This was carried out by fitting the source-EEG envelope FC, after filtering the signal in the different EEG frequency bands, and downsampling to a 100 Hz sampling rate.

For the sleep EEG-fMRI dataset (**Figure 7**) [78], we swept the coupling parameter, *K*, from 0 to 3 (41 values in total), and the target value for ISP, *ρ*, from 2 to 3 (41 values as well), fixing *r*^*α*^ = 0.5 and *τ* = 2 sec. We simulated 660 seconds and discarded the first 60 sec for stabilization. We used the pyramidal neurons’ firing rates to simulate fMRI-like signals through the Balloon-Windkessel model [48], averaging FCs across the 50 simulation seeds. We used firing rates for both α and γ subpopulations scaled by *r*^*α*^ The SC matrix came from the connectome of the young participants dataset [11].

### 4.7 Model fitting

For EEG and fMRI FC matrices, we maximized the SSIM [84], used to compare empirical and simulated FC matrices in whole-brain simulations [76].

For the simultaneous EEG and fMRI dataset, after running the model, we aimed to fit jointly the EEG power spectrum and fMRI FC. First, we calculated the relative power of each EEG band using the Welch method, generating a spectral vector of three entries (relative θ, α, and β power) for empirical and simulated data. Then, we compared the distance between the empirical and simulated relative power using the Clarkson Distance (say, x is the simulated power vector and y the empirical power vector):

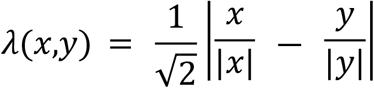

where |*x*| is the Euclidean norm of x. We took 1 - *λ*(*x,y*) as the goodness of fit between spectral vectors *x,y*.

Finally, we calculated the distance between empirical and simulated spectral vectors for each parameter combination and further analyzed only those that with a goodness of fit ≥ 0.85. From this subset of parameters, we maximized the SSIM between empirical and simulated FCs.

### 4.8 Bifurcation analysis

Bifurcation analysis was performed using the XPP-Auto software, and the figures were made using custom code in Python.

### 4.9 Statistical analysis

Given that statistical p-values could be artificially inflated by sample size in computer simulations (varying the number of seeds), instead of statistical tests, we report results using Cohen’s D for effect size. Usually, Cohen’s D is interpreted as very small (|D| < 0.2), small (0.2 < |D| < 0.5), moderate (0.5 < |D| < 0.8), large (0.8 < |D| < 1.2), very large (1.2 < |D| < 2) [85].

### 4.10 Code and data availability

All the codes and basic data used to perform the simulations are freely available in GitHub at https://github.com/carlosmig/EEG-Dementias (“JR ReDLat + Sleep” folder), and mirrored in Zenodo (DOI: 10.5281/zenodo.16929492). We used the BrainNet Viewer toolbox for visualization [86]. Raw data are available on request from Drs. Agustín Ibáñez and Natalia Kowalczyk-Grębska. A formal data-sharing agreement is required.

## Acknowledgments

AI is supported by grants from the Multi-partner consortium to expand dementia research in Latin America [ReDLat, supported by Fogarty International Center (FIC), National Institutes of Health, National Institutes of Aging (R01s AG075775, AG057234, AG082056 and AG083799, CARDS-NIH 75N95022C00031), Alzheimer’s Association (SG-20-725707), Rainwater Charitable Foundation – The Bluefield project to cure FTD, and Global Brain Health Institute)], ANID/FONDECYT Regular (1250091, 1210195, 1210176, and 1220995); ANID/PIA/ANILLOS ACT210096; FONDEF ID20I10152, and ANID/FONDAP 15150012. JC is supported by ANID (FONDECYT Postdoctorado #3240042; FONDECYT de Exploración #13240170). This research was supported by the National Science Centre (Poland) Grant: 2013/11/N/HS6/01335, in the years 2013–2017 (to NK-G). The contents of this publication are solely the authors’ responsibility and do not represent the official views of these institutions. RH was partially supported by the Ramón y Cajal Fellowship (RYC2022-035106-I) from FSE/Agencia Estatal de Investigación (AEI), Spanish Ministry of Science and Innovation, and the María de Maeztu Program for units of Excellence in R&D, grant CEX2021-001164-M/10.13039/501100011033.

## Declaration of interest

None to declare

## Contributions

**Conceptualization**: CC-O, PO, AI. **Data curation**: NK-G, RG-G. **Formal analysis**: CC-O, FL, PO, **Funding acquisition**: AI. **Investigation**: CC-O, PO, AI. **Methodology**: CC-O, FL, AI. **Project administration**: PO, AI. **Resources**: PO, AI. **Software**: CC-O. **Supervision**: PO, AI. **Validation**: CC-O, FL, PO, AI. **Visualization**: CC-O, FL, PO. **Writing – original draft**: CC-O, FL, PO, AI. **Writing – review & editing**: CC-O, FL, RH, IM, MG, NK-G, VM, JC, ET, RG-G, HH, PP, PO, AI.

## Notes

### Competing Interest Statement

The authors have declared no competing interest.

